# The contribution of common genetic risk variants for ADHD to a general factor of childhood psychopathology

**DOI:** 10.1101/193573

**Authors:** Isabell Brikell, Henrik Larsson, Yi Lu, Erik Pettersson, Qi Chen, Ralf Kuja-Halkola, Robert Karlsson, Benjamin B Lahey, Paul Lichtenstein, Joanna Martin

**Affiliations:** Department of Medical Epidemiology and Biostatistics, Karolinska Institutet, Stockholm, Sweden; School of Medical Sciences, Örebro University, Örebro, Sweden; Statistical Genetics, Genetics and Computational Biology Department, QIMR Berghofer Medical Research Institute, Brisbane, QLD, Australia; Department of Public Health Sciences, University of Chicago, Chicago, IL, USA; MRC Centre for Neuropsychiatric Genetics and Genomics, Cardiff University, Cardiff, UK

## Abstract

Attention-deficit/hyperactivity disorder (ADHD) is a heritable neurodevelopmental disorder, with common genetic risk variants implicated in the clinical diagnosis and symptoms of ADHD. However, given evidence of comorbidity and genetic overlap across neurodevelopmental and externalizing conditions, it remains unclear whether these genetic risk variants are ADHD-specific. The aim of this study was to evaluate the associations between ADHD genetic risks and related neurodevelopmental and externalizing conditions, and to quantify the extent to which any such associations can be attributed to a general genetic liability towards psychopathology. We derived ADHD polygenic risk scores (PRS) for 13,460 children aged 9 and 12 years from the Child and Adolescent Twin Study in Sweden, using results from an independent meta-analysis of genome-wide association studies of ADHD diagnosis and symptoms. Associations between ADHD PRS, a latent general psychopathology factor, and six latent neurodevelopmental and externalizing factors were estimated using structural equation modelling. ADHD PRS were statistically significantly associated with elevated levels of inattention, hyperactivity/impulsivity, autistic traits, learning difficulties, oppositional-defiant, and conduct problems (standardized regression coefficients=0.07-0.12). Only the association with specific hyperactivity/impulsivity remained significant after accounting for a general psychopathology factor, on which all symptoms loaded positively (standardized mean loading=0.61, range=0.32-0.91). ADHD PRS simultaneously explained 1% (*p-*value<0.001) of the variance in the general psychopathology factor and 0.50% (*p-*value<0.001) in the specific hyperactivity/impulsivity factor. Our results suggest that common genetic risk variants associated with ADHD have largely general pleiotropic effects on neurodevelopmental and externalizing traits in the general population, in addition to a specific association with hyperactivity/impulsivity symptoms.

## Introduction

Attention-deficit/hyperactivity disorder (ADHD) is a heritable neurodevelopmental disorder characterized by inattentive and hyperactive/impulsive symptoms, affecting 5-10% of children worldwide.^1^ It is well-established that genetic factors contribute to ADHD liability. Twin and sibling studies estimate the heritability of ADHD at 70-80%,^2–6^ with similar heritability across symptom dimensions of inattention and hyperactivity/impulsivity^7^ and across diagnosed ADHD and sub-threshold ADHD symptoms in the population.^8, 9^ Recently, the largest GWAS of clinically diagnosed ADHD to date (20,183 cases, 35,191 controls) identified the first 12 genome-wide significant loci for ADHD, and estimated the proportion of phenotypic variance explained by measured single nucleotide polymorphisms (SNPs), known as SNP-heritability, at h^2^_snp_=0.22(standard error[SE]=0.01).^10^ Significant SNP-heritability has also been reported by the largest GWAS meta-analysis of ADHD symptoms in the population to date (N=17,666), with estimates ranging from h^2^_snp_=0.05(SE=0.06) to h^2^_snp_=0.34(SE=0.17).^11^ Notably, the genetic correlation between the GWAS of clinically diagnosed ADHD and the GWAS of ADHD symptoms was estimated at r_g_=0.94 (SE=0.20), suggesting a near complete common variant genetic overlap across clinically defined ADHD and population variation in ADHD symptoms.^10, 11^ Further, polygenic risk scores (PRS) for clinically diagnosed ADHD (i.e., individual scores of the estimated total burden of risk alleles associated with a phenotype)^12^ have been associated with ADHD population symptoms^13, 14^ and with developmental trajectories of ADHD symptoms.^15^ Conversely, PRS for ADHD symptoms have been associated with clinical ADHD.^16^ Together, these findings suggest that clinically diagnosed ADHD and continuously distributed population symptoms are underpinned by a similar genetic architecture on the level of common variants.^8, 10^

Although numerous studies provide strong support for the contribution of common genetic variants to ADHD, it is also well known that ADHD is highly comorbid with other psychiatric disorders and traits. Therefore, one important question is the degree to which such genetic risk factors are ADHD-specific. Findings from population-based twin studies have suggested a moderate to substantial genetic overlap across ADHD and several other neurodevelopmental and externalizing traits, including autistic traits,^17–20^ learning disabilities,^21–23^ oppositional defiant and conduct problems.^24–27^ Further, twin studies have also shown that a latent shared genetic factor can account for up to 45% of co-variance across childhood externalizing, internalizing and phobia symptoms^28, 29^ and 31% of co-variance in childhood neurodevelopmental symptoms.^30^ Similar results have been reported for register-based clinical diagnoses, with one study showing that a general genetic factor explained 10-36% of disorder liability across several psychiatric diagnoses.^31^ Based on such findings it has been suggested that comorbidity across psychiatric disorder and traits may be attributed to a general psychopathology factor underpinned by common genetic variants.^29, 32^ Two studies have assessed the contribution of measured genetic variants for a general psychopathology dimension; Pappa et al (2015) reported a significant SNP-heritability (h^2^_snp_=0.18, SE=0.10) for maternal ratings of total problems on the Child Behavior Checklist (CBCL), which measures internalizing, externalizing and attention problems.^33^ Similarly, Neumann et al (2016) reported a 38% SNP heritability (h^2^_snp_=0.38,SE=0.16) for a general psychopathology factor derived from childhood psychopathology symptoms assessed by multiple raters.^34^ These findings suggest that the co-occurrence of neurodevelopmental, externalizing and emotional symptoms in childhood is, at least in part, due to shared common risk variants.^34^ Some genomic studies that focused specifically on ADHD have indeed reported a significant, but not complete, genetic overlap with related neurodevelopmental^10, 35–37^ and externalizing conditions^38^, whereas other studies have found no such effects.^10, 39–43^ ADHD PRS have been associated with lower levels of cognitive abilities, IQ and poorer educational attainment.^10, 35, 36^ Two studies have reported significant associations between ADHD PRS and autistic-like social-communication traits^37, 44^, whereas other studies found no association with autistic-like traits^39^, nor with clinically-diagnosed ASD.^39–43^ Higher ADHD PRS have been reported in children diagnosed with ADHD and comorbid conduct, relative to children with ADHD-only and controls.^38^ It therefore remains unclear to what extent genetic risk factors associated with ADHD are condition-specific, or if they also influence a broader range of symptoms including neurodevelopmental, externalizing, and more general psychopathology dimensions. The aims of the current study were to: 1) Examine whether ADHD PRS are associated with a range of other neurodevelopmental and externalizing traits in a large general population sample, and 2) Quantify the extent to which any observed associations between ADHD PRS and the aforementioned symptom dimensions can be attributed to a latent general psychopathology factor.

## Methods

### Study population

The Child and Adolescent Twin Study in Sweden (CATSS) is an ongoing twin study targeting all 9-year-old (born after June 1995) and 12-year-old (born before July 1995) twins born in Sweden since July 1992. Parents were contacted for a telephone interview on the twins’ 9^th^ or 12^th^ birthdays. CATSS was approved by the ethics committee at Karolinska Institutet and all participants gave informed consent. The study has been described in detail elsewhere.^45^

### Phenotypic measures

ADHD and related psychiatric symptoms were assessed using the Autism-Tics, ADHD, and Other Comorbidities inventory (A-TAC). A-TAC is a validated questionnaire including 96 items corresponding to DSM-IV definitions of childhood psychiatric symptoms. Questions addressing lifetime symptoms are assessed in relation to same-age peers and include three response categories: “no” (coded 0), “yes, to some extent” (coded 0.5), and “yes” (coded 1). ^45^ We focused on the 49 items measuring inattention, hyperactivity/impulsivity, autistic traits, learning difficulties, oppositional defiant problems and conduct problems.

### Genotyping and imputation in CATSS

A total of 11,551 CATSS twins were genotyped using the Illumina Infinium PsychArray-24 BeadChip. Prior to analysis, stringent quality control (QC) procedures were performed on the genotyped markers and individuals using standardized procedures (see Supplementary Materials and Figure 1). After QC, 561,187 genotyped SNPs and 11,081 samples were retained. Genotypes for another 2,495 MZ twins were imputed from their genotyped co-twin. Genotype imputation was performed in Minimac3^46^ for 13,576 CATSS samples on autosomes using 1000-Genomes data (Phase 3, Version.5) as the reference panel.^47^ We excluded individuals with parent-reported cerebral palsy, Down syndrome, brain injury and chromosomal abnormalities. The final sample with genotype and phenotype data consisted of 13,460 individuals (50% female). To account for population stratification, principal components (PCs) were derived in CATSS using PC analysis in PLINK after LD-pruning and removing genotyped SNPs located in long-range LD regions. We calculated PCs on unrelated individuals and then projected the PCs onto the relatives.

**Figure 1.**
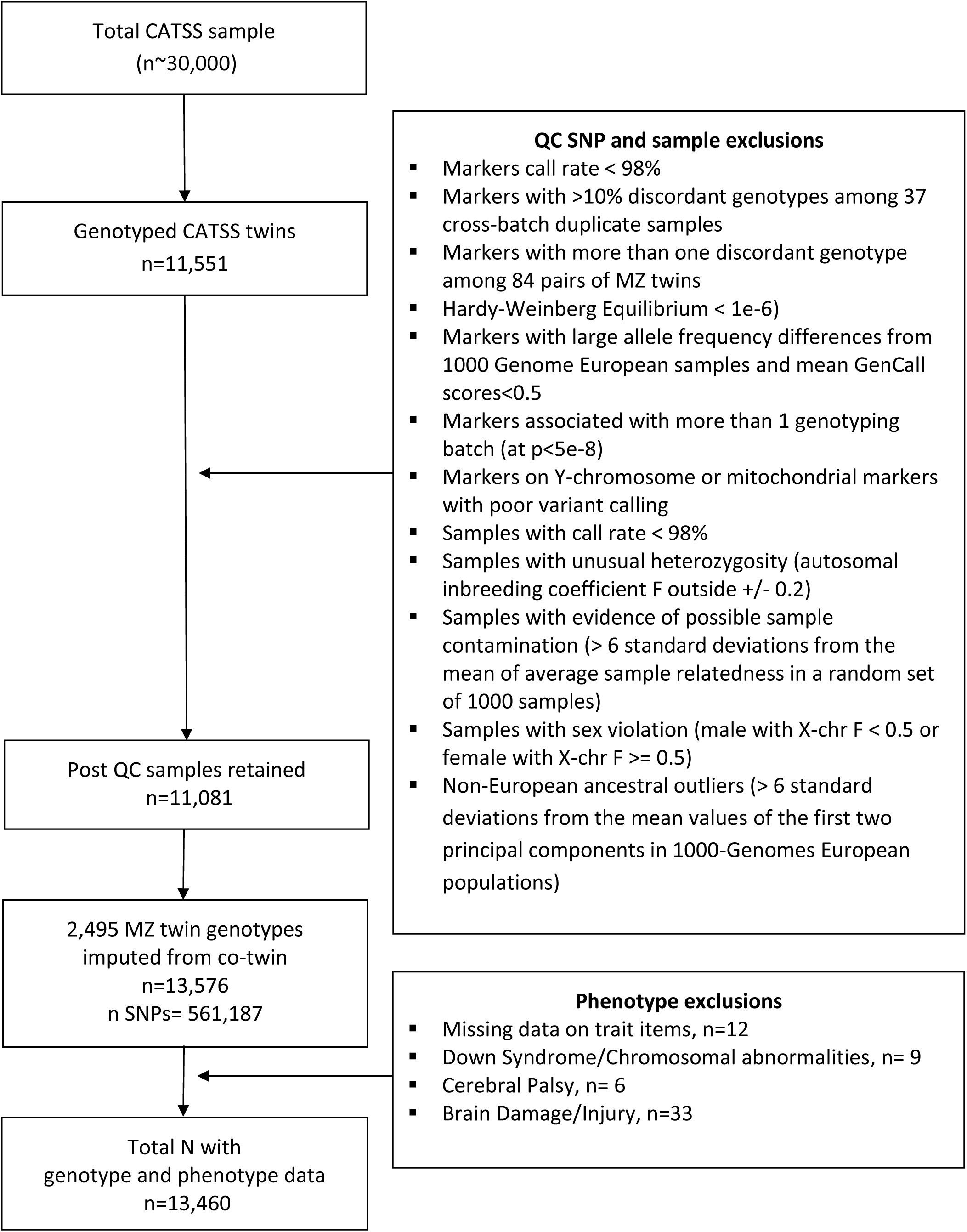
Flow chart of quality control protocol and population selection in the CATSS sample

**Figure 2.**
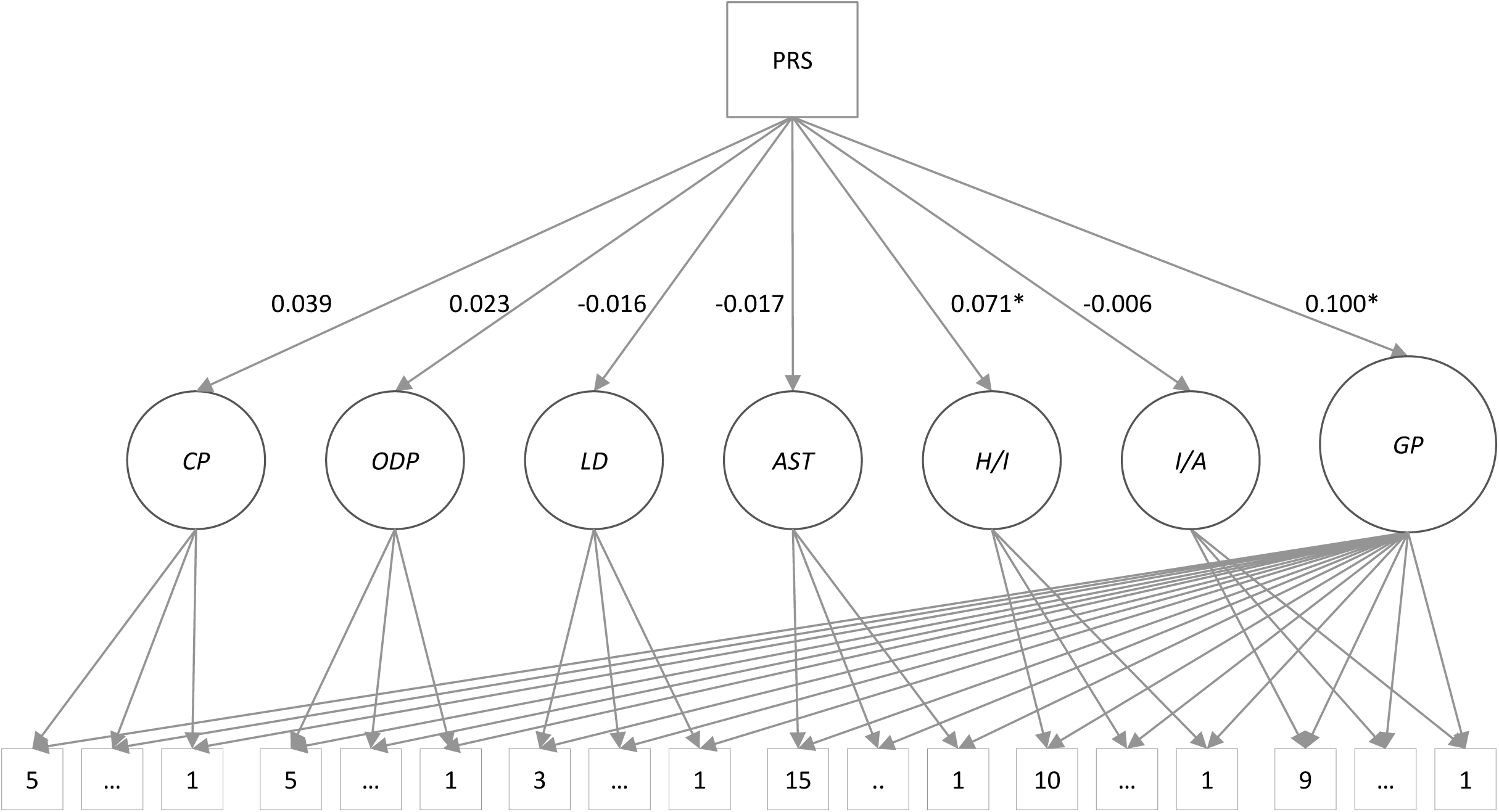
General Psychopathology Factor Model and ADHD PRS associations Latent factor are the depicted as circles. The general psychopathology factor model consisted of a latent general factor (GP) and six latent trait factors reflecting specific inattention (IA), hyperactivity/impulsivity (HI), autistic traits (AST), learning difficulties (LD), oppositional defiant problems (ODP) and conduct problems (CP). Measured variables are depicted as squares; the ADHD PRS derived from the clinical+population ADHD GWAS and the 49 A-TAC symptoms items, with 1…X indicating the number of symptoms loading onto each latent trait variable. For clarity, covariates (age, sex and six principal components), correlations across latent trait factors and path loadings from latent factors onto each measured item are omitted in the above graphical representation. A full graphical representation of the model is presented in the supplementary materials (Figure S2). Variances for all latent factors were fixed at 1. Presented beta estimates are standardized. * Significant at p-value threshold < 0.0001.

### Polygenic risk scores

Polygenic risk scores (PRS) were generated in CATSS based on summary statistics from a meta-analysis of the two largest GWAS of ADHD available to date. The GWAS of clinically diagnosed ADHD (henceforth referred to as the ‘clinical ADHD GWAS’) included 20,183 cases and 35,191 controls and was conducted by The Lundbeck Foundation Initiative for Integrative Psychiatric Research (iPSYCH)^48^ and the Psychiatric Genomics Consortium (PGC) ADHD working group.^10^ The GWAS of ADHD symptoms (henceforth to as the ‘population ADHD GWAS’) included 17,666 children and was conducted by the EArly Genetics and Lifecourse Epidemiology Consortium (EAGLE).^11^ Meta-analysis of the two GWAS (henceforth referred to as the ‘clinical+population ADHD GWAS’) was conducted using a new method, relying on modified sample size-based weights to account for the respective heritabilities, genetic correlation, and measurement scale of each GWAS; See supplementary materials in Demontis et al (2017) for details.^10^ The clinical+population ADHD GWAS not only replicated the 12 genome-wide significant loci identified in the clinical ADHD GWAS, but also identified an additional three loci. The genetic correlation between the two GWAS was 0.94 and there was no evidence of genome-wide significant heterogeneity across the samples, indicating strong genetic overlap^10^. These results suggest that the clinical+population ADHD GWAS should provide the most powerful discovery sample available to derive ADHD PRS. We calculated standardized betas for each SNP, based on available information of z-scores, effective sample size and allele frequency in the clinical+population ADHD GWAS, in line with recommendations.^49^ ADHD PRS were derived in CATSS from best-guess imputed genotypes across a range of *p*-value thresholds (0.00001≤*P*_*T*_≤1). Indels, multi-allelic and symmetric/ambiguous SNPs were excluded. Autosomal SNPs with a minor allele frequency (MAF)≥0.05 and good imputation quality (INFO score)≥0.8 were clumped (linkage disequilibrium threshold *R*^2^>0.1, ±1000 kb) using PLINK.v.1.9.^50^ Retained reference alleles were scored across the set of SNPs in PLINK (applying the command --score no-mean-imputation) using standard procedures.^51, 52^ In line with previous publications, we used the PRS including SNPs at a threshold of *P*_*T*_≤0.50 for the main analysis^15, 36^. We also derived ADHD PRS from the clinical ADHD GWAS and the population ADHD GWAS for sensitivity analyses.

### Statistical Analyses

We estimated the associations between ADHD PRS and inattention, hyperactivity/impulsivity, autistic traits, learning difficulties, oppositional defiant and conduct problems using two measurement models identified via confirmatory factor analysis (CFA). We first fitted a correlated factor model to the 49 A-TAC symptoms; items in each scale were set to load onto a single latent factor, which were allowed to correlate with each other (Supplementary Figure S1).

Second, we fitted a general psychopathology factor model (Supplementary Figure S2), also referred to as a bi-factor model. In addition to the aforementioned six latent trait factors, this model included a general psychopathology factor on which all 49 symptoms loaded. The general factor is assumed to account for the covariance among the traits, and the specific latent trait factors for the unique variance in each trait.^53^ The six latent trait factors were allowed to correlate, whereas the correlations between the general factor and the latent trait factors were constrained at 0.

In both models, the latent factors were regressed on ADHD PRS using structural equation modelling (SEM), with sex, age and the first six PCs (to account for possible population stratification) included as covariates. We compared model fit using the root mean square error of approximation (RMSEA), the comparative fit index (CFI), and the chi-square difference test.^54^ To account for the non-independence of twin data, family clusters were specified and standard errors were estimated using as and wich estimator. Analyses were run using Mplus.^55^

### Sensitivity Analyses

We performed several sensitivity analyses to test the robustness of our results. First, we tested whether ADHD PRS showed similar associations with the latent factors across a range of p-value thresholds (0.00001≤*P*_*T*_≤1). Second, to test whether observed associations were driven by ADHD cases, we re-ran analyses excluding children with an ICD diagnosis of ADHD (obtained from the Swedish Patient Register) or an A-TAC based DSM diagnosis of ADHD (defined as eight or more DSM-based inattention and/or hyperactivity/impulsivity symptoms). Third, we stratified the analyses by sex. Fourth, we excluded one MZ twin per pair to ensure estimates were not inflated by the inclusion of genetically identical individuals. Finally, we re-ran the analysis using ADHD PRS derived separately from the clinical ADHD GWAS and the population ADHD GWAS.

## Results

### Correlated Factor Model and ADHD PRS associations

The correlated factor model fit the data well (CFI=0.94; RMSEA=0.03, 95%CI=0.03–0.03; χ^2^=14076.72, df=1112). All symptoms loaded positively and significantly onto their corresponding latent trait factor. Factor loadings are reported in Table 1. The six neurodevelopmental and externalizing trait factors were strongly and positively correlated (mean *r*=0.68 range=0.42-0.82, *p*<0.0001) (Supplementary Figure S3). Standardized regression results are reported in Table 2; higher ADHD PRS were significantly (*p*<0.0001) associated with higher symptom levels in each latent trait factor, after adjusting for covariates (Inattention β=.10; Hyperactivity/Impulsivity β=.12; Autistic traits β=.07; Learning difficulties β=.07; Oppositional defiant problems β=.08; Conduct problems β=.10).

**Table 1.**
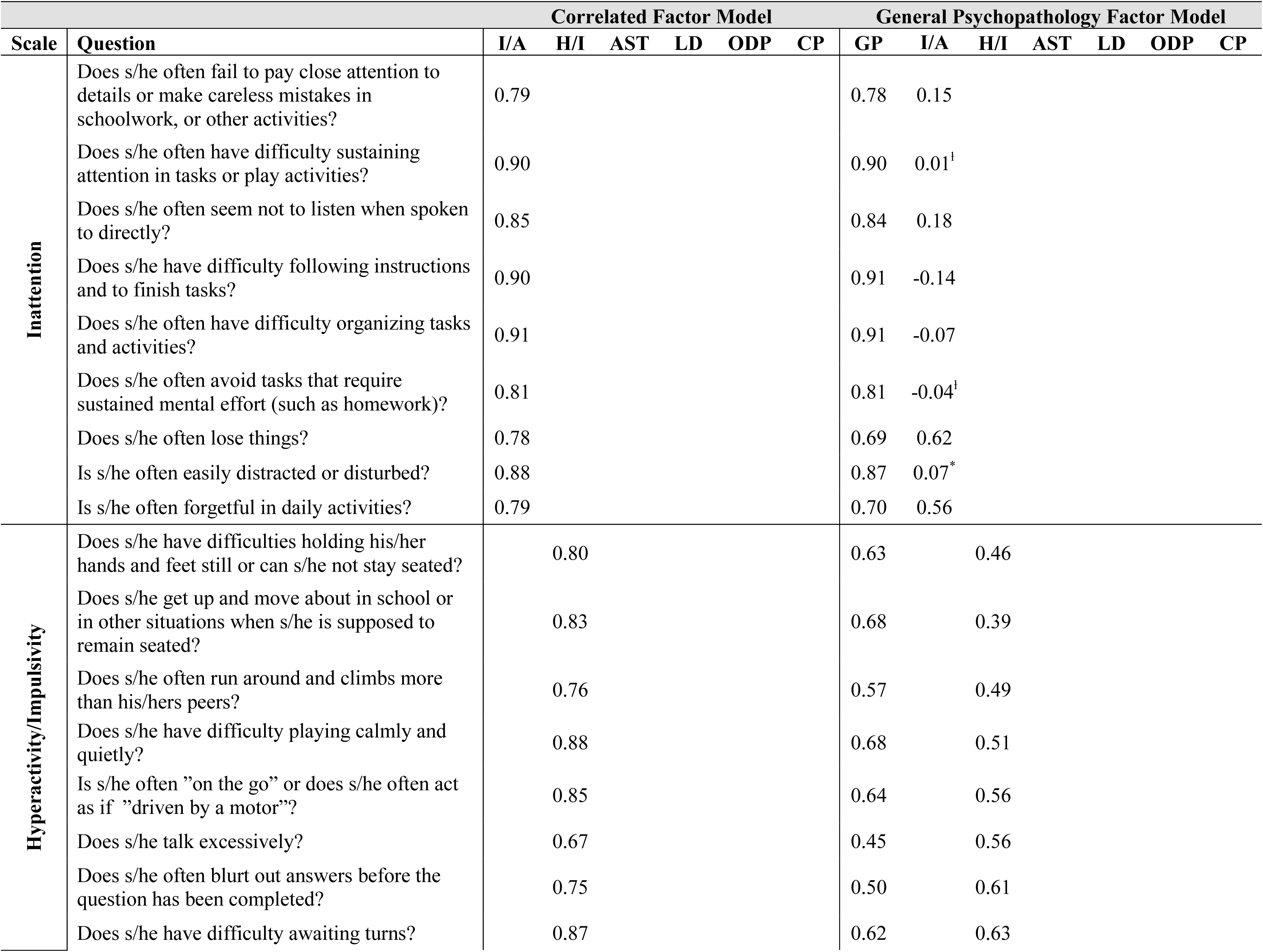
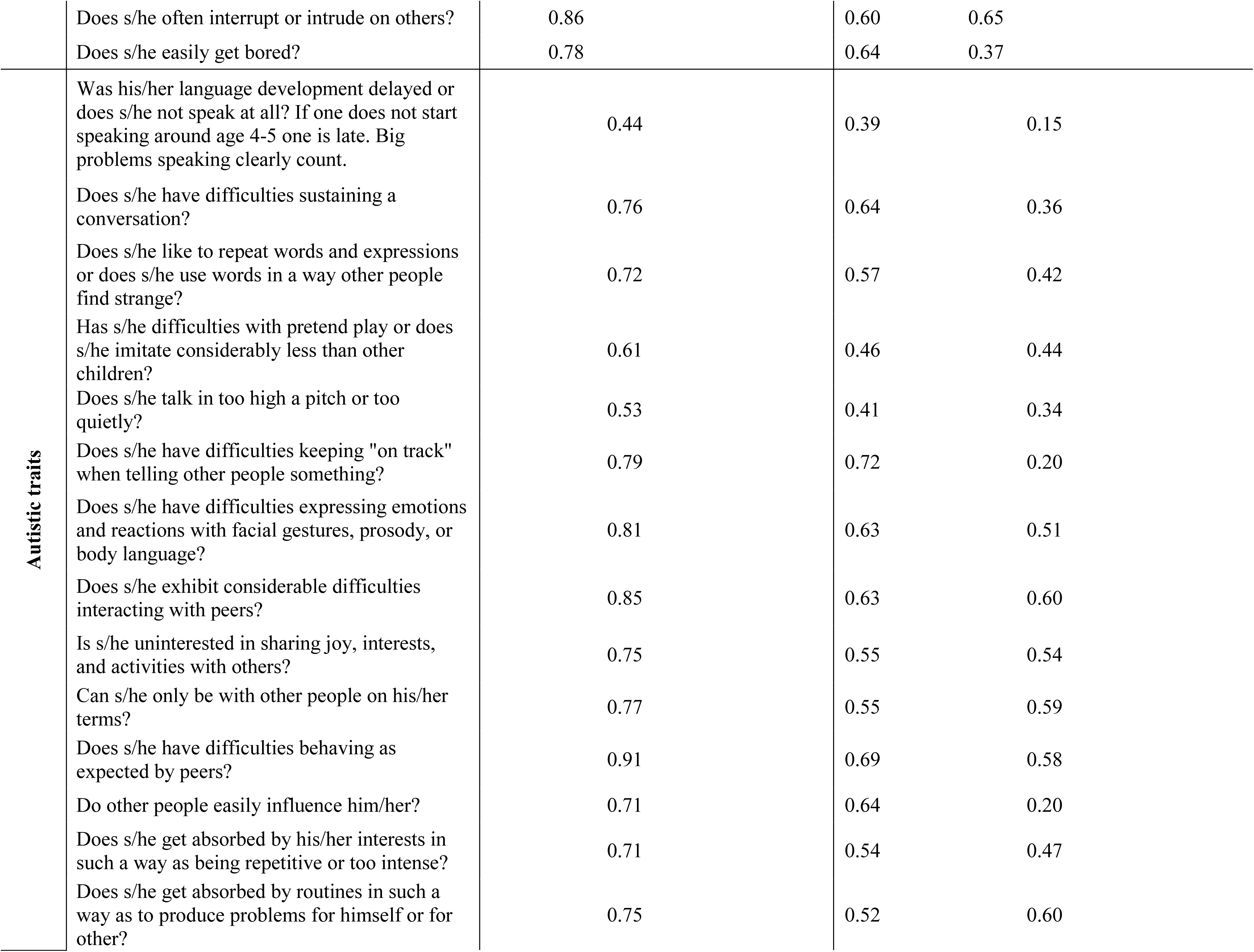
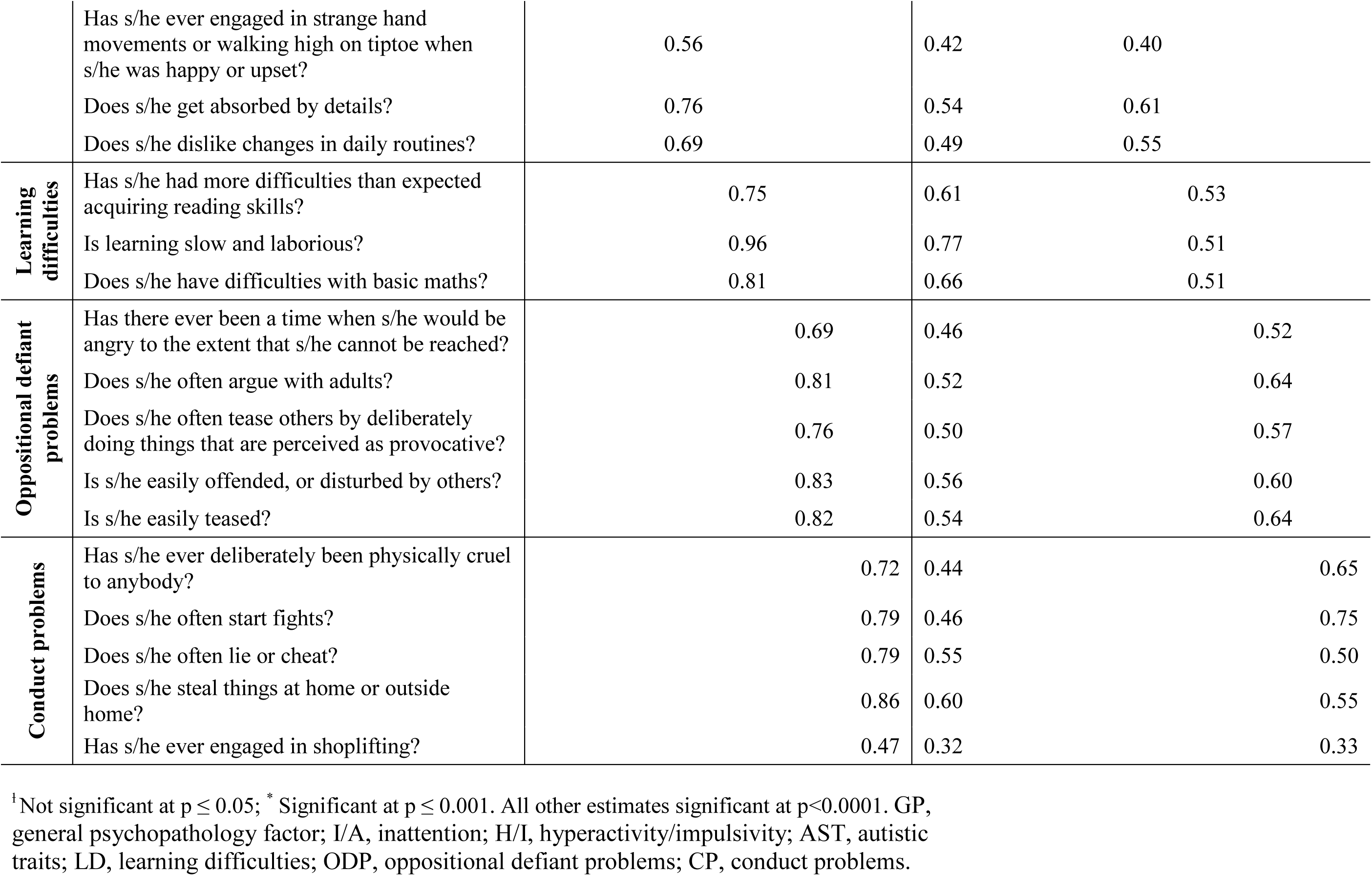
Standardized factor loadings from the Correlated Factor Model and the General Psychopathology Factor Model

**Table 2.** Association between ADHD PRS and latent trait factors in the Correlated Factor Model and the General Psychopathology Factor model

| Latent Factor | Correlated Factor Model |  |  |  | General Factor Model |  |  |  |
| --- | --- | --- | --- | --- | --- | --- | --- | --- |
|  | Beta | SE | p | % variance explained | Beta | SE | p | % variance explained |
| GP | - | - | - |  | 0.10 | 0.01 | <.0001 | 1.00% |
| I/A | 0.10 | 0.01 | <.0001 | 0.96% | -0.01 | 0.02 | 0.718 | 0.00% |
| H/I | 0.12 | 0.01 | <.0001 | 1.44% | 0.07 | 0.01 | <.0001 | 0.50% |
| AST | 0.07 | 0.01 | <.0001 | 0.45% | -0.02 | 0.02 | 0.314 | 0.03% |
| LD | 0.07 | 0.02 | <.0001 | 0.49% | -0.02 | 0.02 | 0.408 | 0.03% |
| ODP | 0.08 | 0.02 | <.0001 | 0.69% | 0.02 | 0.02 | 0.154 | 0.05% |
| CP | 0.10 | 0.02 | <.0001 | 0.94% | 0.04 | 0.03 | 0.115 | 0.15% |

Reported results reflect the association with ADHD PRS derived from the clinical+population ADHD GWAS meta-analysis. The correlated factors model included six latent trait factors and no G factor; the general psychopathology factor model included a general factor model that loaded on all items, in addition to the six correlated latent trait factors. Models were adjusted for sex, age and six principal components. Reported betas are standardized. % variance explained is $\beta^2$ .
GP, general psychopathology factor; I/A, inattention; H/I, hyperactivity/impulsivity; AST, autistic traits; LD, learning difficulties; ODP, oppositional defiant problems; CP, conduct problems.

### General Psychopathology Factor Model and ADHD PRS associations

The general psychopathology factor model also fit the data well (CFI=0.96; RMSEA=0.02, 95%CI=0.02–0.03; χ^2^=9358.46, df=1063). Furthermore, omitting the general factor resulted in a statistically significant decrease in model fit based on the Chi-square difference test (Δχ^2^=2893.16, Δdf=48, *p*<.0001). All symptoms loaded positively and significantly onto the general psychopathology factor (standardized mean loading=0.61, mean SE=0.01, range=0.32-0.91). The general factor explained 62% of the covariance across traits (explained common variance [ECV]).^56^ Loadings on the general factor were strongest for symptoms of inattention (standardized mean loading =0.82) and learning difficulties (standardized mean loading=0.68), and weakest for oppositional defiant (standardized mean loading =0.59) and conduct problems (standardized mean loading=0.56). Notably, only two inattentive symptoms continued to load onto the specific inattentive factor after accounting for the general factor (Table 1). Several of the correlations across the specific factors, for example between inattention and autistic traits, were attenuated or even became negative after accounting for the general psychopathology factor (Supplementary Figure S3).

Standardized regression results are reported in Table 2; higher ADHD PRS were significantly associated with higher scores on the general psychopathology factor (β=.10, *p*<0.001), explaining 1% of the variance in the general factor after adjusting for covariates. After accounting for the general factor, only the association between ADHD PRS and the specific hyperactivity/impulsivity factor remained significant (β=.07, *p*<0.001), explaining 0.50% of the of the variance in specific hyperactivity/impulsivity after adjusting for covariates.

### Sensitivity Analyses

ADHD PRS showed similar associations across the range of p-value thresholds (Supplementary Figures 4 and 5), with one exception; whereas there was no significant association (*p*=0.115) between ADHD PRS (*P*_*T*_≤0.50) and conduct problems in the main analyses, significant associations (*p*<.05) were observed across three more conservative p-value thresholds (*P*_*T*_≤0.00001; *P*_*T*_≤0.05; *P*_*T*_≤0.1). In the sensitivity analysis excluding 864 ADHD cases, ADHD PRS remained statistically significantly associated with the general factor and specific hyperactivity/impulsivity (Supplementary Table S1). Analyses in males showed the same pattern of results as in the whole sample, although the variance explained was larger (general factor:1.54%; hyperactivity/impulsivity: 0.61%). In contrast, the variance explained was lower among females (general factor:0.53%; hyperactivity/impulsivity:0.34%). Results excluding one MZ twin per pair did not differ markedly from the results in the full cohort (Supplementary Table S1). Results using ADHD PRS derived from the clinical ADHD GWAS were nearly identical to the results using ADHD PRS from the clinical+population ADHD GWAS. In contrast, results based on PRS derived from the population ADHD GWAS were substantially attenuated (Supplementary Table S2).

## Discussion

Results from this study show that common genetic variants which increase the risk for ADHD are not only associated with symptom dimensions of ADHD traits in the general population, but also with several related neurodevelopmental and externalizing traits. ADHD PRS were significantly associated with symptoms of inattention, hyperactivity/impulsivity, autistic traits, learning difficulties, oppositional defiant problems, and conduct problems. Importantly, when modelling the shared variance across these traits using a general psychopathology factor, we found that the associations were largely accounted for by a general liability towards childhood psychopathology. The significant association between ADHD PRS and a general childhood psychopathology factor suggests that common SNPs associated with ADHD in GWAS have pleiotropic effects across symptom dimensions of neurodevelopmental and externalizing behaviors in children in the general population.

Beyond the association between ADHD PRS and a general psychopathology factor, we found that ADHD PRS also showed a unique association with specific hyperactivity/impulsivity symptoms. About 2/3 of the association between ADHD PRS and hyperactivity/impulsivity could be attributed to general variance shared across childhood neurodevelopmental and externalizing traits, and about 1/3 to variance specific to hyperactivity/impulsivity. There are several plausible interpretations for the genetic association with specific hyperactivity/impulsivity. Hyperactivity/impulsivity symptoms are more strongly linked to aggressive and externalizing behaviors than inattentive symtoms^24^, and ADHD with comorbid conduct disorder has been found to show a greater genetic load as measured by ADHD PRS.^38^ Although not significant in our main analyses, specific conduct problems did show significant associations with ADHD PRS across three more conservative *p*-value thresholds, even after accounting for a general psychopathology factor. Although this effect would need to be confirmed in future studies, one possible explanation is therefore that ADHD PRS captures genetic risk variants uniquely or more strongly associated with hyperactive/impulsive and aggressive behaviors, over and above more general pleotropic effects on childhood psychopathology. Another possibility is that hyperactive/impulsive symptoms are stronger drivers of ADHD diagnosis, thus leading to an over-representation of combined and primarily hyperactive/impulsive ADHD in the discovery sample of clinically diagnosed ADHD. In contrast, there was no significant association between ADHD PRS and specific inattentive symptoms, after accounting for cross-trait covariance via the general psychopathology factor. This lack of association is likely explained by the fact that the majority of inattentive symptoms loaded very strongly onto the general factor, and the specific inattention factor only captured variance in two items, largely reflecting forgetfulness.

In the correlated factor model, we found a significant association between ADHD PRS and autistic traits. Similar results have previously been found in some^37, 44^, but not all studies^39^. Differences across studies may be due to the power of previously-available discovery samples, the size of target samples, and/or measurement differences. However, our results suggest that this association was not trait-specific, but rather due to a general genetic liability towards childhood psychopathology. This highlights the need to, whenever possible, account for the inter-related nature of psychiatric disorders and traits when studying their genetic architecture.^34^ Further, our results provide additional evidence for the contribution of measured genetic variants, specifically those related to risk of ADHD, to a general psychopathology factor, which is in line with recent studies reporting significant SNP heritability (18-38%) of a general psychopathology factor in childhood.^33, 34^

Analyses stratified by sex showed a somewhat stronger association in males relative to females, particularly for the associations between ADHD PRS, the general psychopathology factor and specific hyperactivity/impulsivity. Whilst this could reflect sex-specific genetic differences, it is more likely to be due to lower levels of hyperactivity/impulsivity in females,^57^ potential under-reporting of neurodevelopmental and externalizing traits in females by parents and/or fewer female cases in the clinical ADHD GWAS discovery data (∼25%).^10^

Beyond demonstrating the importance of a shared general genetic liability for neurodevelopmental and externalizing traits, our results also provide additional support for the dimensional view of ADHD. Several previous studies using diverse methodology (i.e. twin methods, PRS and linkage-disequilibrium score regression) have shown that genetic factors associated with clinically diagnosed ADHD are also linked to population variation of ADHD symptoms.^8, 10, 11, 13, 14, 16^ We extend these findings by showing that PRS based on a meta-analysis of GWAS of clinical diagnosis and population symptoms of ADHD is also linked to a range of childhood psychopathology traits in the general population. Associations were very similar when using ADHD PRS derived from the clinical GWAS only, suggesting that the observed associations were largely driven by SNP effects identified in the clinical ADHD GWAS.^10^ This is not surprising as the study population in the clinical ADHD GWAS^10^ was considerably larger than in the population ADHD GWAS.^11^ Nonetheless, our results indicate that combining summary statistics from GWAS of clinically-diagnosed ADHD and ADHD symptoms is a valid approach and that the identified genetic risk variants are also associated with related childhood psychopathology symptoms in the population. Considering the large sample sizes required for detection of genetic loci implicated in complex traits and disorders, the possibility to pool genetic data across psychiatric dimensions and across clinical and population-based samples in order to increase statistical power is very attractive. Although such efforts pose challenges in terms of sample and measurement heterogeneity,^58^ our results suggest that recently developed multivariate methods for joint GWAS analysis of related disorders and traits are important not only to boost power, but also to identify genetic variants with pleiotropic effects across psychopathology.^58–60^

### Limitations

Results from this study must be interpreted in the context of the study limitations. Firstly, we only had parent ratings of childhood neurodevelopmental and externalizing traits, which can lead to informant bias when estimating the general psychopathology factor.^34^ Although we cannot exclude the possibility that parent ratings may have inflated the cross-trait covariance, the pattern and strength of the general factor model in our study did not differ greatly from studies using multiple informants^34^ or register-based clinical diagnoses.^31^ Secondly, we only included neurodevelopmental and externalizing traits, as both are early onset psychopathology dimensions that are strongly linked to ADHD. Nonetheless, there is also evidence that ADHD in childhood is linked to internalizing traits^28, 61^ and genomic studies have reported genetic associations across clinical ADHD and depression.^10, 43^ Future studies assessing genetic specificity and generality in relation to ADHD should therefore aim to also include internalizing traits. Third, our study is cross-sectional. Although there is strong evidence for genetic stability in ADHD,^44, 62^ regulation of gene expression is a dynamic process, such that age-specific effects and developmental trajectories may influence the strength and nature of cross-disorder genetic associations.^15, 44^

### Conclusion

Results from this study indicate that majority of genetic risk variants implicated in ADHD represent a general genetic liability towards childhood psychopathology, reflecting a multi-dimensional continuum of genetic risk that underpins both neurodevelopmental processes and externalizing behaviors. Beyond contributing to shared genetic liability, ADHD PRS also seemed to capture genetic risk with stronger and/or unique effects on specific hyperactivity/impulsivity symptoms. Our findings suggest that adopting a more general, multivariate framework, considering the intersection between neurodevelopmental and externalizing childhood traits and disorders is crucial when studying the genetic architecture of childhood psychopathology. Further, our findings also suggest that combining data across psychopathology dimensions and across clinical and population samples may increase the power to identify genetic risk related to childhood psychopathology in future GWAS.

## Acknowledgments

We gratefully acknowledge the contribution of the participants in the Child and Adolescent Twin Study in Sweden (CATSS) and their families. CATSS is supported by the Swedish Council for Working Life (no 2012-1678 and 2014-0834), funds under the ALF agreement (no 2014-0322) and the Swedish Research Council (no 340-2013-5867 and 2014-3831). Dr. Martin was supported by the Wellcome Trust (Grant No: 106047). The funders had no role in study design, data collection and analysis, decision to publish or preparation of the manuscript. We also acknowledge and thank Dr Patrick Magnusson for his work and expertise on the genotyping, imputation and quality control process of the genetic samples in CATSS, and Dr Raymond Walters for his statistical advice and expertise on the use of the iPSYCH/PGC+EAGLE GWAS meta-analysis discovery data.

## Conflicts of Interest

Ms. Brikell, Drs.Yi, Pettersson, Chen, Kuja-Halkola, Karlsson, Lahey, and Martin declare no potential conflicts of interest. Dr. Larsson has served as a speaker for Eli-Lilly and Shire and has received research grants from Shire. Dr. Lichtenstein has served as a speaker for Medice. All outside the submitted work.

